# Altered effective connectivity in sensorimotor cortices: a novel signature of severity and clinical course in depression

**DOI:** 10.1101/2021.03.14.435324

**Authors:** Dipanjan Ray, Dmitry Bezmaternykh, Mikhail Mel’nikov, Karl J Friston, Moumita Das

**Author notes:** D.R., and M.D. conceived the present project. D.B., and M.M. performed experiments, and collected data. D.R., and M.D. performed data analysis. M.D., and K.J.F. supervised the project. D.R., and M.D. wrote the manuscript. D.B., M.M., and K.J.F. edited the manuscript. The authors declare no competing interest.

## Abstract

Functional neuroimaging research on depression has traditionally targeted neural networks associated with the psychological aspects of depression. In this study, instead, we focus on alterations of sensorimotor function in depression. We used resting-state functional MRI data and Dynamic Causal Modeling (DCM) to assess the hypothesis that depression is associated with aberrant effective connectivity within and between key regions in the sensorimotor hierarchy. Using hierarchical modeling of between-subject effects in DCM with Parametric Empirical Bayes we first established the architecture of effective connectivity in sensorimotor cortices. We found that in (interoceptive and exteroceptive) sensory cortices across participants, the backward connections are predominantly inhibitory whereas the forward connections are mainly excitatory in nature. In motor cortices these parities were reversed. With increasing depression severity, these patterns are depreciated in exteroceptive and motor cortices and augmented in the interoceptive cortex: an observation that speaks to depressive symptomatology. We established the robustness of these results in a leave-one-out cross validation analysis and by reproducing the main results in a follow-up dataset. Interestingly, with (non-pharmacological) treatment, depression associated changes in backward and forward effective connectivity partially reverted to group mean levels. Overall, altered effective connectivity in sensorimotor cortices emerges as a promising and quantifiable candidate marker of depression severity and treatment response.

**Significance Statement:** Research into neurobiology of depression primarily focuses on its complex psychological aspects. Here, we propose an alternative approach and target sensorimotor alterations - a prominent but often neglected feature of depression. We demonstrated using resting-state fMRI data and computational modelling that top-down and bottom-up information flow in sensory and motor cortices is altered with increasing depression severity in a way that is consistent with depression symptoms. Depression associated changes were found to be consistent across sessions, amenable to treatment and of effect size sufficiently large to predict whether somebody has mild or severe depression. These results pave the way for a new avenue of research into the neural underpinnings of mental health conditions.

---

The search for the neurological bases of depression has provided many important insights, yet we are far from a comprehensive, translatable understanding (1–4). This warrants further research and, possibly, new approaches.

Neuroimaging research on depression largely focuses on complex affective and psychological components of depression, the prefrontal cortex and limbic formation being two of the most investigated brain regions (5). At the network level, apart from the fronto-limbic circuitry, default mode network, cognitive control network, and corticostriatal circuits are some of the major neurocircuits that are known to be involved in depression (6–19).

However, depression is an embodied phenomenon and is known to cause alterations in several sensorimotor functions. Persons suffering from depression, for example, are known to have reduced visual contrast sensitivity (20), impaired auditory processing of non-speech stimuli (21), and increased pain tolerance for exteroceptive stimulation (22). In addition to these exteroceptive alterations, depression has been shown to cause interoceptive changes like decreased pain tolerance for interoceptive stimulation (22) and reduced heartbeat perception accuracy (23). The psychomotor retardation (reduced speed, slow speaking rate, delayed motor initiation, body immobility, loss of facial expression (24)) is a prominent feature of depression. Indeed, psychomotor retardation has been played an important role in the descriptive characterization of depression and melancholia since their nosological inception (24–29). Darwin (30) described overt psychomotor symptoms in sad people who “no longer wish for action but remain motionless and passive, or may occasionally rock themselves to and fro”. In the following decades, scholars such as Emil Kraepelin developed the concept further and established its clinical utility (25, 26). Among later researchers, Carl Wernicke (31), Karl Kleist (32) and Karl Leonhard (33) contributed to our refined understanding of psychomotor abnormalities. Lastly, rumination, an important feature of depression (34), has prominent sensorimotor components.

Although there are a few neuroimaging studies of sensorimotor changes in depression, our understanding of sensory and motor function of brain is undergoing a paradigm shift. Spear-headed by predictive coding and related theoretical frame-works, there is an emerging consensus among neuroscientists that perception is not a simple ‘bottom-up’ mechanism of progressive abstraction of sensory input (35–37). Bottom-up, top-down and intrinsic neuronal message passing play distinct but crucial roles. This general idea is also applicable to motor function (see active inference (38)). Motivated by these novel insights, we analysed effective connectivity (spectral dynamic causal modelling (39)) in resting state functional MRI data among hierarchical sensorimotor regions in unmedicated depression patients and neurotypical individuals. For exteroceptive perception, effective connectivity among the lateral frontal pole - one of the terminal regions of sensory relays - and primary visual, auditory, and somatosensory cortices was considered. Effective connectivity between anterior and posterior insula was characterised for interoception and between supplementary motor area and primary motor cortex was analysed for motor function (Figure 4). Both group mean effective connectivity and connections showing significant association with Beck Depression Inventory (BDI) scores (40) (after controlling for age and sex) were identified. In a leave-one-out cross-validation (41) - using parametric empirical Bayesian - the effect size was estimated. A subset of participants, who were either treated with cognitive behaviour therapy (42), neurofeedback therapy (43) or not treated were scanned again a few months later and same analysis was implemented, with the addition of treatment effect as a covariate.

## Results

### The primary experiment

#### Accuracy of DCM model estimation

The accuracy of DCM estimates of effective connectivity for individual participants was excellent. Across participants, the minimum percentage variance-explained by DCM - when fitted to the observed (cross spectra) data - were 73.55%, 68.84%, and 55.00% for left motor, exteroceptive, and interoceptive networks, respectively. For right hemisphere ROIs, these values were 63.2%, 50.79%, and 30.75%. In general, for most participants variance explained was 80% or more.

#### Effective connectivity

Results are displayed in Figure 1 and detailed further in supplementary Figure 1

**Fig. 1.**
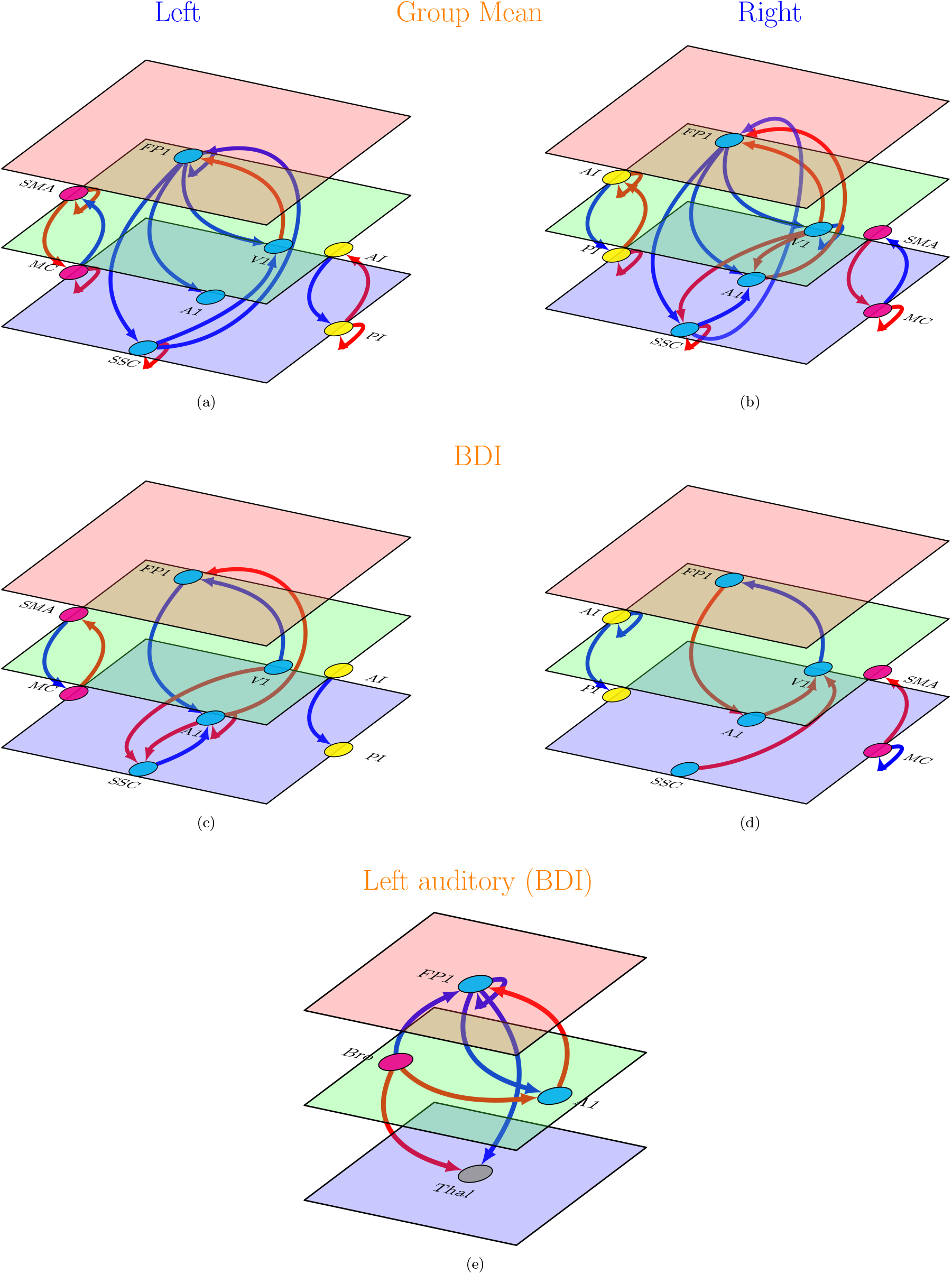
Effective connectivity in the primary study (left and right hemispheres). (a),(b): Group mean effective connectivity in sensory and motor networks. Arrow colours code nature of connections red, excitatory; blue, inhibitory. (c),(d): Connections showing significant association with Beck depression inventory (BDI) scores in sensory and motor networks. Arrow colours code direction of connectivity changes relative to the group mean: red, increased; blue, decreased. (e): Connections showing significant association with Beck depression inventory (BDI) scores in a network composed of left thalamus, left primary auditory cortex, Broca’s region and left lateral frontal pole. For all subfigures line thickness is kept constant and does not code for the effect size. For the exact values of the estimated connectivity parameters see supplementary Figure 1. Colours of the planes denote position of the node in cortical hierarchy. Green is higher than blue, red is higher than both blue and green. SMA: supplementary motor area, MC: primary motor cortex, FP1: lateral frontal pole, V1: primary visual cortex, A1: primary auditory cortex, SSC: primary somatosensory cortex, AI: anterior insula, PI: posterior insula. Bro: Broca’s region. Thal: Left thalamus. The images were created using tikz-network (https://github.com/hackl/tikz-network) package in LATEX.

### Group mean effective connectivity

The mean effective connectivity among sensorimotor regions is depicted in Figure 1 (a) and (b). Among extensive network of connections in both hemispheres,the most consistent pattern emerged in the forward and backward effective connectivity. In sensory regions (exteroceptive and interoceptive), backward connections were inhibitory, whereas forward connections were excitatory (exception: SSC to FP1 connection). In motor regions, opposite was true (backward: excitatory, forward: inhibitory).

### Changes in effective connectivity with BDI scores

The connections that showed an association with BDI scores are shown in Figure 1 (c) and (d). As with mean connectivity, the severity associated changes were most consistent in (extrinsic or between region) forward and backward connections across both hemispheres. For exteroceptive and motor cortices, with increasing BDI scores top-down and bottom-up effective connectivity show changes in the opposite direction with respect to group level estimation. For example, in exteroceptive sensory regions (with one exception, see below) bottom-up connections become more negative and top-down connections become more positive (i.e., disinhibition). In motor regions, top-down connections become more negative and bottom-up connections become more positive. In interoceptive regions top-down inhibitory influences are enhanced.

### Effective connectivity analysis for left auditory regions

One notable exception to general pattern of changes in exteroceptive sensory regions with BDI scores was found in left auditory regions. Here top-down inhibitory and bottom-up excitatory influences were enhanced with depression. One possible explanation is that this effect reflects enhanced rumination and self-speech in depression (please note that the left auditory cortex is specialized for speech perception). To further probe this hypothesis we implemented spectral DCM analysis among left thalamus, Broca’s area, left A1, and left FP1 regions. We found that left A1 was driven mainly by Broca’s area rather than the left Thalamus (see second sub-figure below). We will return to this observation in discussion.

### Cross Validation

In a leave-one-out cross-validation, among all six networks, the left exteroceptive network was found to predict BDI scores at a significant level of *α* = 0.05 (see Table 1). When individual connections were considered, three connections of left exteroceptive network, namely left V1 to FP1 (corr=0.23, p-value=0.036), left A1 to SSC (corr=0.22, p-value=0.045), left SSC to A1 (corr=0.23, p-value=0.03) were found to have significant predictive power for BDI scores. Note that these measures of effect size correspond to out of sample measures (i.e., the effect sizes one would see using effective connectivity estimates from new participants).

**Table 1.**
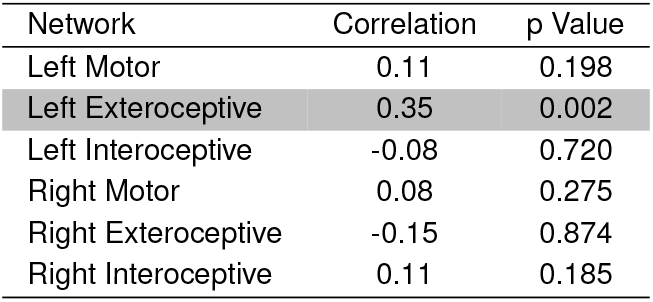
Leave-one-out cross validation: results from the primary study.

### The follow-up experiment

#### Accuracy of DCM model estimation

As in primary analyses, the accuracy of DCM predictions for individual participants was excellent for the follow up study. The minimum percentage variance-explained by DCM model estimation across participants were 57.14%, 76.90%, and 73.33% for left motor, exteroceptive, and interoceptive networks and 76.02%, 68.70%, and 44.06% for right motor, exteroceptive, and interoceptive networks. For most of the participants variance explained was 80% or more.

#### Change in BDI scores

The BDI scores of participants during the first and the second sessions are plotted in Figure 2. As evident from the figure, for most of the participants in the treatment as well as no treatment group, BDI scores improved with time; however, improvement was more prominent in the treatment group. This was also corroborated by statistical testing. The paired samples Wilcoxon test indicated that BDI scores during the first session were statistically significantly higher than the second session for both groups at significance level *α* = 0.05. However, at significance level *α* = 0.01, this held true only for the treatment group (p-value = 0.009491) but not for the no treatment group (p-value = 0.01176).

**Fig. 2.**
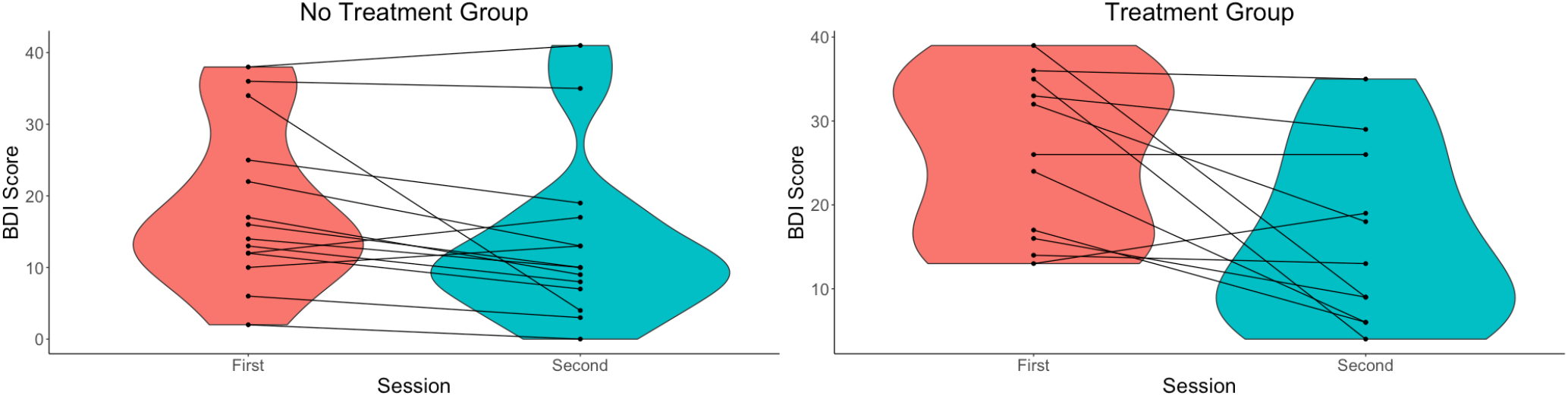
Violin plots of the Beck depression inventory (BDI) scores in (a) no treatment and (b) treatment groups across sessions. A violin plot is a box plot with the width of the box proportional to the estimated density of the observed data.

#### Effective connectivity

Results are displayed in Figure 3 and are further detailed in supplementary Figure 2.

**Fig. 3.**
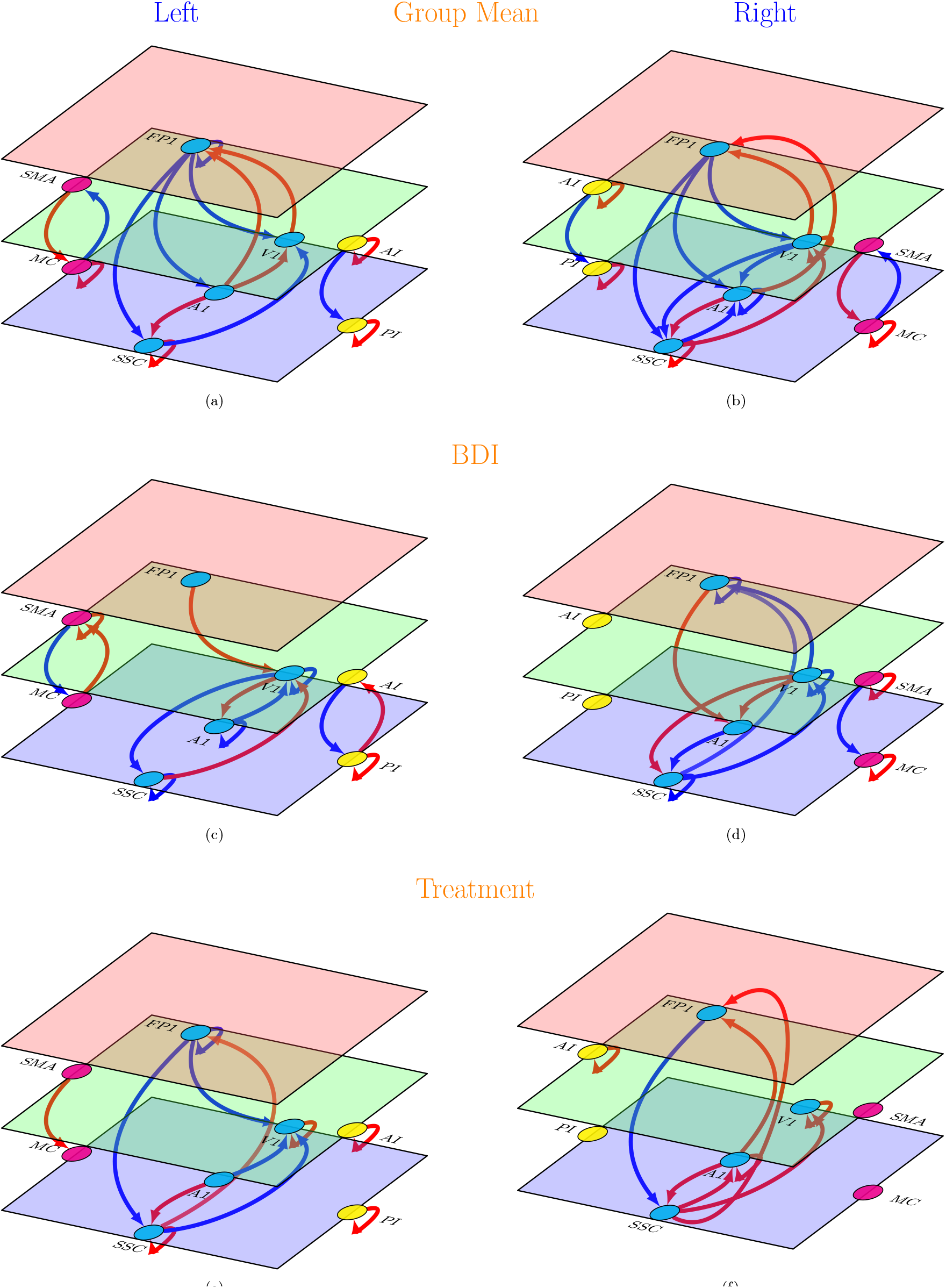
Effective connectivity in the follow-up study (left and right hemispheres). (a),(b): Group mean effective connectivity. Arrow colours code nature of connections red, excitatory; blue, inhibitory. (c),(d): Connections showing significant association with Beck depression inventory (BDI) scores. Arrow colours code direction of connectivity changes relative to the group mean: red, increased; blue, decreased. (e),(f): Connections showing significant association with treatment (treatment vs no treatment). Arrow colours code direction of connectivity changes relative to the group mean: red, increased; blue, decreased. For all subfigures line thickness is kept constant and does not code for the effect size. For the exact values of the estimated connectivity parameters see supplementary Figure 2. Colours of the planes denote position of the node in cortical hierarchy. Green is higher than blue, red is higher than both blue and green. SMA: supplementary motor area, MC: primary motor cortex, FP1: lateral frontal pole, V1: primary visual cortex, A1: primary auditory cortex, SSC: primary somatosensory cortex, AI: anterior insula, PI: posterior insula. The images were created using tikz-network (https://github.com/hackl/tikz-network) package in LATEX.

### Group mean effective connectivity

Overall, the main pattern of mean effective connectivity was reproduced by the follow up analysis. The backward connections in exteroceptive and interoceptive cortices are inhibitory and forward connections are excitatory. The opposite pattern was observed in bilateral motor cortices.

### Changes in effective connectivity with BDI scores

Like mean effective connectivity, the changes in effective connectivity between hierarchical cortical regions with increasing depression severity follow the same pattern found in the primary analysis: with increasing BDI scores the top-down and bottom-up mean effective connectivity is enhanced in the interoceptive network and is diminished in exteroceptive and motor networks.

### Changes in effective connectivity with treatment

With treatment, top-down and bottom-up effective connectivity revert towards group mean levels, i.e., in the exteroceptive network, top-down effective connections become more inhibitory and bottom-up connections becomes more excitatory; whereas in the motor network top-down connections became more excitatory. In the interoceptive network, no change in top-down or bottom-up effective connectivity survived at the 95% threshold set for the posterior probability of the estimated parameters.

### Cross Validation

In a leave-one-out cross-validation, none of the effective connections were found to predict BDI scores at a significant level of *α* = 0.05 (see Table 2).

**Table 2.**
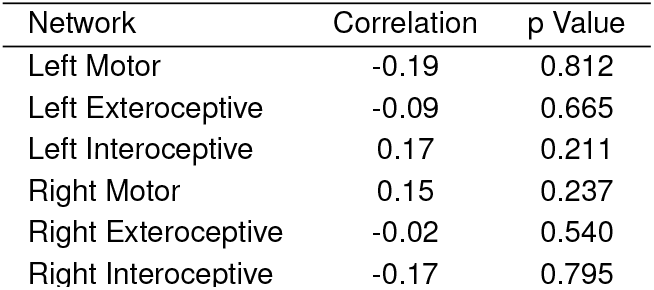
leave-one-out cross validation: results from the follow-up study.

## Discussion

Overall, the most exciting findings from our study are the average backward (top-down) and forward (bottom-up) effective connectivity in sensory and motor cortices that showed consistent patterns across hemispheres and sessions and consistent changes with depression severity and treatment. The backward effective connections in exteroceptive and interoceptive sensory networks were predominantly inhibitory in nature while forward connections were predominantly excitatory (except SSC to FP1 connections in primary experiment). The opposite pattern was observed in bilateral motor networks. With increased depression scores, this pattern is weakened in exteroceptive and motor networks and is strengthened in the interoceptive network. Interestingly, with treatment, a partial recovery towards the group average was observed. In leave-one-out cross validation analysis, connections in left exteroceptive networks were found to have sufficiently large effect size to predict whether somebody has a high or a low BDI score.

There is a growing recognition that the depression is associated with dysfunction of distributed brain networks rather than of individual brain regions (44, 45). Four networks have been the focus of most of the published research in this area: the affective network (AN), reward network (RN), default mode network (DMN), and cognitive control network (CCN). Hyperconnectivity among the regions of AN (12, 14) and DMN (6, 10, 13, 16, 46) has been consistently reported in depression. Enhanced resting state functional connectivity in AN and DMN has been postulated to be associated with negative affectivity and maladaptive rumination in depression patients. Hypoconnectivity in RN (7, 17, 18) and CCN (9, 11, 47) has been another consistent finding in depression (but also see (8, 48) for divergent findings). Anhedonia and ineffective cognitive control over emotional processing seen in depression have been attributed to diminished interactions among the regions of RN and CCN, respectively.

As evident from above, the affective and psychological components of depression have been the prime focus of neurobiological research on depression. Yet, several sensorimotor interventions including light, music, tone, physical exercise are well known to modulate mood and depressive symptoms (49). Association of depression with visual (50, 51) or hearing impairment (52–54) is also well established. Depression, in turn, gives rise to several sensorimotor alterations. Some of them, for instance, psychomotor retardation or agitation and feelings of fatigue are part of the diagnostic criteria for depression (55). Besides, there is a repertoire of subjective feelings that depressed patients experience. These include pain in several parts of the body, chest discomfort, feeling cold or nauseous, heaviness of limbs, feeling of emptiness, to mention a few (56). These feelings change the subjective experience of one’s own body and one’s sense of relatedness with the world outside.

There are only a few neuroimaging studies that independently examined functional connectivity in sensory and motor networks as biomarkers for depression. Among them, one recent study (57) found reduced within and between-network functional connectivity in auditory and visual networks associated with depression. In another study, Kang et al. (58) demonstrated that the primary somatosensory area-thalamic functional connectivity is abnormal in major depressive disorder. Moreno-Ortega et al. (59) showed that including resting state functional connectivity within the visual network in the analysis greatly increases the predictive power for the treatment response to electroconvulsive therapy in depression compared to model consisting of only AN and DMN.

However, our understanding of neuronal mechanisms underlying sensory perception is going through a major shift. There is an emerging consensus that perception is not a passive ‘bottom-up’ mechanism of progressive abstraction from sensory input and both bottom-up and top-down connectivity between hierarchically organized brain regions play crucial roles in perception. This recognition has led to several theoretical frameworks highlighting the importance of top-down information flow in the context of sensory perception. The most prominent of them - predictive coding (35–37) - has also been extended to motor function (see active inference (38)). These novel insights motivated us to analyse effective connectivity among hierarchical brain regions in sensory and motor cortices. In contrast to data-driven approaches (e.g., functional connectivity analyses) mentioned above, ours is a model-based approach informed by theoretical frameworks and empirical knowledge of functional architectures. In motor regions we chose primary motor area and supplementary motor area. The later is responsible for planning complex movements of the contralateral extremities and is posited to occupy a higher level of hierarchy in the motor system. Similarly, in interoceptive cortex we chose posterior and anterior insula based on known role of the insula in interoception and a posterior to anterior hierarchical organization in the insula (60, 61). For exteroception, we selected three primary sensory cortices: visual, auditory, and somatosensory and the lateral frontal pole - the terminal relay station for exteroceptive sensory information (62, 63).

A consistent and intriguing finding from our study is top-down inhibitory and bottom-up excitatory average effective connectivity in sensory cortices; a pattern that reverses in motor cortices. The pattern in sensory cortices is consistent with the role of top-down predictions explaining away prediction errors at lower levels, via interactions with inhibitory interneurons in canonical microcircuits (as proposed by the predictive coding framework). In other words, although long-range connections in the brain are excitatory (i.e., glutamatergic), backward connections may preferentially target inhibitory interneurons in superficial and deep layers to evince an overall decrease in neuronal message passing. In predictive coding, this is often read as ‘explaining away’ prediction errors at lower levels in sensory cortical hierarchies (64). However, the completely opposite pattern was observed in the motor network. Descending excitatory connections in the motor system may reflect one of two things. First, it could be a reflection of the fact that ascending prediction errors in the executive motor system may play a small role – because these prediction errors are thought to be resolved through cortical spinal reflexes; i.e., through action (38). Put simply, in sensory hierarchies exteroceptive prediction errors are caused by bottom-up sensory input, which are resolved by (inhibitory) top-down predictions. Conversely, in motor hierarchies prediction errors are generated by (excitatory) top-down proprioceptive predictions, which are resolved by motor reflexes at the level of the spinal-cord. An alternative explanation is that descending predictions include predictions of precision that may mediate things like attention and sensory attenuation (65–67). In this instance, there can be an explaining away of certain prediction errors, while there precision may be increased, resulting in an overall excitatory drive. In other words, some descending predictions may be of proprioceptive gain that mediates the selection of intended movements. In this context it is noteworthy that descending predictions of precision play an important role in active inference accounts of psychiatric conditions – in which the synaptic pathophysiology and psychopathology can be accounted for by a failure of sensory attenuation; namely, the attenuation or suspension of the precision of sensory prediction errors. This failure of attention and attenuation has been used to explain several conditions, including autism, schizophrenia, Parkinson’s disease and depression (68–72). The current results are particularly prescient in relation to formulations of depression and mood disorder in terms of active inference; namely, how actions are selected by inferring ‘what to do next’. Clark, Watson and Friston (71) review the evidence for depression as a computational pathology in the proprioceptive and interoceptive (behavioural and autonomic) domain. They conclude “emotional states reflect the precision associated with neurobiological predictions over interoceptive states”. The current results are consistent with this formulation but draw special attention to proprioceptive predictions in the sensorimotor system. In this setting, the attenuation of descending effective connectivity – to the executive motor cortex with increasing depression severity – is consistent with a failure to deploy sensorimotor precision appropriately during action selection. In turn, this is consistent with a failure to form precise (subpersonal) beliefs about ‘what to do next’, at higher levels in the sensorimotor hierarchy. An extreme example of the ensuing psychomotor poverty may be the bradykinesia of Parkinson’s disease, which has a clear neuromodulatory (dopaminergic) aetiology. Please see (73) for further discussion.

In line with the marked consistency of the patterns of average effective connectivity - across hemispheres and sessions - the changes in effective connectivity with depression severity were also conserved across sessions and corroborate well with depressive symptomatology. Instead of categorically dividing participants into patients and neurotypical subjects, we examined (across participants) variation of effective connectivity with depression severity as assessed by the Beck Depression Inventory. This leverages the heterogeneity within each group that might contain useful clinical information (74). With increasing depression severity, the patterns found in top-down and bottom-up connections at the group level are weakened in exteroceptive (except the left auditory cortex-see below) and motor cortices and strengthened in the interoceptive cortex. Depreciation in exteroceptive networks is in line with he reduced visual contrast sensitivity (20) and impaired auditory processing of non-speech stimuli (21). Psychomotor poverty or retardation is a prominent feature of depression 24) that might well be reflected in the weakening of motor network effective connectivity. The enhancement in the interoceptive network is consistent with increased interoceptive e.g., pain) sensitivity (22) in depression. On the contrary, a ew studies reported a subtle but non-significant association of depression with decreased interoceptive awareness like reduced heartbeat perception accuracy (75, 76). However, small sample sizes and/or inclusion of individuals with mild or comorbid presentations of depression may undermine this claim 77, 78). Moreover, Pollatos,Traut-Mattausch, Eva and Rainer 23) found that a negative relationship between depression and heartbeat perception accuracy is only present in those with relatively higher trait anxiety. Thus, it might reflect an interaction of anxiety with depression. Furthermore, Dunn, Dalgleish, Ogilvie and Lawrence (79) found that heartbeat perception accuracy was affected in mild depression but, paradoxically, was not affected in more severely depressed group thus further complicating the association.

One notable exception - to general pattern of changes in ffective connectivity within exteroceptive network with BDI cores - was found in left auditory regions. Here top-down nhibitory and bottom-up excitatory influences were enhanced with depression. One possible explanation is that this reflects enhanced rumination and self-speech in depression; noting hat left auditory cortex is specialized for speech perception 80). Rumination is implicated in the development, severity and maintenance of depression and other psychiatric disorders 81–83). Given the central role of rumination in depression, t has been considered a key target in modern cognitive and behavioural therapies (84). One of the most salient features of rumination is that it is mostly expressed in a verbal modality 85–87). In other words, while ruminating, we are mostly alking to ourselves silently. Thus, enhancement of effective onnectivity within auditory network, with increasing BDI cores, might reflect depressive rumination during the acquisition of resting-state scans. To further probe this hypothesis we implemented spectral DCM effective connectivity analysis among left thalamus, Broca’s area, left A1 and left FP1 egions. Broca’s area, also known as the left inferior frontal gyrus (LIFG), is involved in production of both outer and inner speech (e.g., (88)). We hypothesized that if the change in the pattern of effective connectivity with increasing depression everity is associated with rumination, left auditory area (A1) would be driven mainly by Broca’s area. Conversely, if it reflects some form of aberrant sensory processing, left thalamus will be main driver of left A1 (89). DCM analysis demonstrated that with increasing BDI score effective connectivity rom left Broca’s area to left A1 becomes more excitatory but here is no significant change in effective connectivity from eft Thalamus to left A1, thus providing an indirect support or the rumination hypothesis. It is noteworthy here that a previously published report of the same data found that the independent component - representing the left auditory network - also included the insular cortex in the depression group but not in the healthy participants. Based on several lesion (90–94) and neuroimaging (95, 96) studies, the left insula has been proposed as a brain region involved in motor control of speech production including pre-articulatory motor responses (97–99). This lends further support to depressive rumination conjecture.

The model comparison discussed above furnishes clear evidence for changes in a number of extrinsic (between region) and intrinsic (within region) connections that underwrite depression, as scored with the BDI. One might ask whether these changes can be used diagnostically in individual patients. In other words, are the underlying effect sizes sufficiently large to predict whether somebody has a high or a low BDI score. This question goes beyond whether there is evidence for an association and addresses the utility of connectivity phenotyping for personalised medicine. One can address this using out of sample estimates of the effect size using cross validation under a parametric empirical Bayesian scheme (41). In other words, one can establish the predictive validity by withholding a particular subject and ask whether one could have predicted the BDI score given the effective connectivity estimates from that subject. This question can be posed at the level of a single perception accuracy was affected in mild depression but, para-connection or sets of connections. For example, when looking at single connections, three connections in the left hemisphere all showed a significant out of sample correlation with BDI score. This suggests that a nontrivial amount of variance in the BDI score could be explained by effective connectivity. This variance explained increased when considering the left exteroceptive network – attaining a correlation coefficient of 0.35 or, an R-squared of about 10% (which was extremely significant *p <* 0.001). Although relatively small from a psychological perspective, this is almost an order of magnitude greater than the variance can be explained by genomic phenotypes (100, 101).

Clinicopathological significance of effective connectivity in sensory and motor cortices is further supported by the DCM analysis of treatment-associated changes in connectivity in the follow up study. Several top-down and bottom-up connections in bilateral exteroceptive and motor cortices were found to be associated with treatment. More importantly, the parity of these connections is opposite to the connections showing an association with depression severity, suggesting a prognostic relevance of these connectivity measures. Remarkably, none of the feedforward or feedback connections in the interoceptive cortex was found to be associated with treatment, but the clinical significance of this finding is unknown. Taken together, the patterned alterations in bidirectional connectivity with BDI scores and treatment offer a strong case for effective connectivity in sensory and motor cortices as a biomarker for depression.

A few words on the computational method used in the current work. DCM was introduced originally to model neuronal responses to external perturbation (e.g., sensory stimulation or task demands). DCM for resting state fMRI was subsequently introduced in Stochastic DCM (102). Stochastic DCMs differ from deterministic DCMs by allowing for physiological noise due to endogenous stochastic fluctuations in neuronal and vascular responses, known technically as system or state-noise.The opportunity to model endogenous (autonomous) fluctuations opened the door to identify the functional architectures (effective connectivity) subtending endogenous fluctuations observed in resting-state studies. A more efficient approach for resting state data was subsequently introduced which is based on fitting observed complex fMRI cross spectra (39) (For more details see Materials and Methods). This later approach, known as spectral DCM, was employed in the present study.

Findings from the current study should be appreciated within the context of certain limitations. Although our study sample was modestly large for neuroimaging measures - and we undertook steps like cross-validation and replication of the main results to ensure the generalizability of our findings - replication in an independent sample would be an important next step. Secondly, in the context of connectivity analysis, there are several potential confounding factors other than age and sex of the participants that we have not controlled for. For example, level of anxiety in individuals could affect top-down information flow in the brain (103). Anxiety is also a common comorbidity found in depression patients (104). None of our participants reported to be diagnosed with anxiety disorders. However, the presence of subclinical anxiety was not ruled out or controlled for. We will consider testing for the association of anxiety with effective connectivity in sensory and motor networks in a companion paper. A third limitation of our study is that the analysis relied solely on BDI scores of depression. There are a large number of rating scales for assessing depression severity: some are observer rating scales, for example the Hamilton Depression Rating Scale (HDRS) and the Montgomery-Åsberg Depression Rating Scale (MADRS), others are self-rating scales (for example BDI). Each scale has its own advantages and limitations (105). Thus, the present neuroimaging findings could be further validated with a combination of observer rating scales and objective behavioural measures of depression (e.g. (106)).

In summary, our results advance our mechanistic understanding of depression pathophysiology. Traditional accounts of depression (e.g. Beck’s (107) cognitive model) have neglected bodily symptoms (79). The present work re-establishes depression as an embodied phenomenon by demonstrating that effective connectivity in sensory and motor cortices affords a promising neural signature of depression. It also establishes the generalizability and predictive validity of this novel marker– and may portend a new avenue of research into the neural underpinnings and therapeutic interventions of depression and other mental health conditions.

## Materials and Methods

### Participant characteristics

Fifty-one adult patients (mean age: 32.78 years, SD: 8.89, 38 females, 13 males) with a diagnosis of mild depressive episode or moderate depressive episode according to ICD-10 and twenty-one adult individuals (mean age: 33.8 years, SD: 8.5, 15 females, 8 males) with no history of neurological or psychiatric illness participated. Depressed participants were either referred by a qualified psychiatrist or invited through advertisement in a popular local newspaper and then assessed by the same psychiatrist. Inclusion criterion were first diagnosed mild or moderate depressive episode and age between 18 and 55 years. Exclusion criteria were: previous depressive episodes, bipolar depression, seasonal depression, depression secondary to other psychiatric or somatic condition, serious risk of suicide, serious neurological and psychiatric comorbidities, alcohol or other substance abuse or dependence, lifetime history of psychotic disorders, contraindications to MRI, extremely impaired vision, IQ score below 70, any psychotropic medication (including antidepressants), and any medication altering blood pressure (that could influence fMRI signal). Healthy participants were volunteers recruited by word of mouth or via advertisement in social networks. Inclusion and exclusion criteria for healthy volunteers were the same, except for the presence of depressive episodes. The depressed and neurotypical participants did not differ in level of intelligence (mean (SD) Raven’s Progressive Matrices test score, for neurotypicals: 105.9(16.5), for depression patients: 103.7(14.6)). All participants gave informed consent in accordance with the Declaration of Helsinki. Ethical review board of Research Institute of Molecular Biology and Biophysics approved the study. Beck depression inventory evaluation could not be done on four patients and three neurotypical participants. Consequently, sixtyfive participants were included in the final analysis.

Twenty-nine depression patients from the primary study were included in the follow-up study (gap between two sessions, minimum: 56 days, maximum: 234 days). Among them fifteen individuals received no treatment, eight received cognitive behavioural therapy (CBT) and six received neurofeedback therapy (NFBT). BDI scores could not be retrieved for one participant during the first scan and for four participants during the second scan and subsequently twenty-four participants were included in the final analysis. We checked for systemic differences between participants who attended both the sessions and who dropped out. A Mann-Whitney test failed to show between-group differences in age, IQ, and emotional variables at a significance level of 0.05. At the same significance level, the chi-square analyses failed to show significant differences between two groups in terms of sex ratio and mild/moderate depression ratio.

It is noteworthy here, data from a subset of participants from the present study has been published (46, 108, 109). However, those works mainly employed a data-driven approach based on independent component analysis (ICA) decomposition of the whole-brain data and correlation based (undirected) functional connectivity analysis unlike the current study that tests a specific hypothesis by investigating (directed) effective connectivity in functionally characterised brain regions.

### Brain MRI acquisition

The fMRI acquisition was carried out in the International Tomography Center, Novosibirsk. Imaging data were acquired with an Ingenia (Philips) 3T scanner using a 32-channel dStream HeadSpine coil (digital). The structural and functional images had the following parameters: Structural MRI: T1 3D TFE, Field of View: 250 × 250 × 280 mm^3^, TR/TE=7.5/3.7 ms, Flip Angle= 8°, Voxel size: 1× 1 × 1 mm^3^. Functional MRI: T2* Single shot SPIR EPI, Field of View: 220 × 220 mm^2^, TR/TE=2500/35 ms, Flip Angle= 90°, Voxel size: 2 × 2 × 5 mm^3^, 25 slices.

During the resting state sequence (duration: four minutes each), participants were instructed to lie still and motionless in the scanner with their eyes closed while letting their mind wander.

### Preprocessing

The pre-processing and statistical analysis of fMRI data were executed with the SPM12 v7771 toolbox (Statistical Parametric Mapping, http://www.fil.ion.ucl.ac.uk/spm). The initial five scans were discarded to allow the magnetization to stabilize to a steady state. Prior to statistical analysis, images were slice-time corrected, realigned with the mean image, motion corrected, coregistered with the corresponding T1-weighted images, normalized to a Montreal Neurological Institute (MNI, https://www.mcgill.ca) reference template and resampled to 4 × 4 × 5 mm^3^. During motion correction, 2nd-degree B-Spline interpolation was us ed for estimation and 4th-degree B-Spline for reslicing. Coregistration used mutual information objective function while normalization used 4th-degree B-Spline interpolation. Images were smoothed with a full-width at half-maximum (FWHM) Gaussian kernel 4 × 4 × 10 mm^3^ and further denoised by regressing out several nuisance signals, including the Friston-24 head motion parameters and signals from cerebrospinal fluid and white matter. Temporal high pass filtering above 1/128 Hz was employed to remove low-frequency drifts caused by physiological and physical (scanner related) noises.

### Spectral Dynamic Causal Modelling and Parametric Empirical Bayes

The Spectral DCM approach using DCM12.5 as implemented in SPM12 v7771 (http://www.fil.ion.ucl.ac.uk/spm) was used to estimate the effective connectivity within each network. Dynamic causal modelling (DCM) is Bayesian framework that infers the causal architecture of distributed neuronal systems from the observable BOLD (blood-oxygen-level-dependent) activity recorded in fMRI. It is primarily based on two equations. First, the neuonal state equation models the change of a neuronal state-vector in ime, depending on modulation of connectivity within a distributed ystem and experimental perturbations. Second, an empirically alidated hemodynamic model that describes the transformation of neuronal state into a BOLD response. For task fMRI, external stimuli usually forms the external perturbation component. For resting-state fMRI, in the absence of external stimuli – a stochastic component capturing neural fluctuations is included in the model nd the neural state equation can be represented as

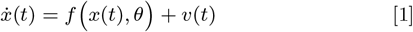

where 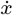 is the rate of change of the neuronal states x, *θ* represents unknown parameters (i.e., intrinsic effective connectivity) and v(t) is the stochastic process modelling the random neuronal fluctuations hat drive the resting-state activity. The observation equation could be written as:

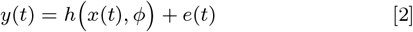

Here, y(t) is the observed BOLD activity, *φ* are the unknown parameters of the (haemodynamic) observation function, and e(t) is the stochastic process representing the measurement or observation noise.

Spectral DCM offers a computationally efficient inversion of the tochastic model for resting state fMRI. Spectral DCM simplifies the enerative model by replacing the original BOLD time-series with heir second-order statistics (i.e., cross spectra). This allows circumenting estimation of time varying fluctuations in neuronal states by estimating their covariance, which is time invariant. In other words, the problem of estimating hidden neuronal states disappears and is replaced by the problem of estimating their correlation functions of time or spectral densities over frequencies (and observation noise) where a scale free (power law) form is used (motivated from previous works on noise in fMRI (110) and underlying neuronal activity (111, 112)) as follows:

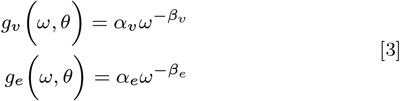

Here, {*α, β*} ⊂ *θ* are the parameters controlling the amplitudes and exponents of the spectral density of the neural fluctuations. Finally, tandard Bayesian model inversion (i.e. Variational Laplace) is used o infer the parameters of the models from the observed signal. A detailed mathematical treatment of spectral DCM can be found in (Ref: 39) and (113).

Time series for DCM analysis were extracted for each region of interest by taking the first principal components of the time series from all voxels included in the masks for that region. Masks were defined according to SPM Anatomy toolbox (114). The regions of interest for each network are depicted in Figure 4. We also adjusted data for “effects of interest”, thus effectively mean-correcting the time series.

**Fig. 4.**
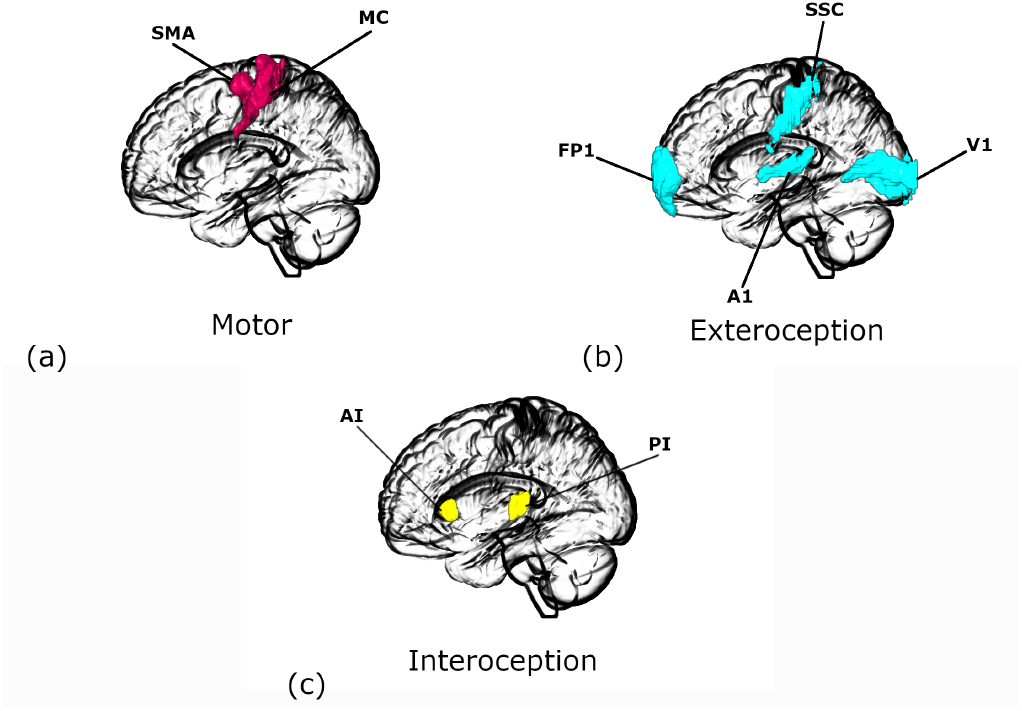
Regions of interest for (a) Motor, (b) Exteroceptive, and (c) Interoceptive networks. SMA: supplementary motor area, MC: primary motor cortex, FP1: lateral frontal pole, V1: primary visual cortex, A1: primary auditory cortex, SSC: primary somatosensory cortex, AI: anterior insula, PI: posterior insula. The images were created using MRIcroGL(https://www.nitrc.org/projects/mricrogl/) program.

At the first level, fully-connected models (i.e., between all nodes plus self-loops) were estimated for each subject individually, separately for bilateral exteroceptive, interoceptive and motor networks.

A basic diagnostic of the success of model inversion is to look at the average percentage variance-explained by DCM model estimation when fitted to the observed (cross spectra) data. We implemented this diagnostic test across participants.

At the second (group) level, we used parametric empirical Bayes (PEB) — a between-subjects hierarchical Bayesian model over parameters — which models how individual (within-subject) connections relate to different between-subjects effects (41, 115)(Friston, Zeidman and Litvak, 2015; Friston et al., 2016). Unlike a classical test (e.g., t-test), it uses the full posterior density over the parameters from each subject’s DCM – both the expected strength of each connection and the associated uncertainty (i.e., posterior covariance) – to inform the group-level result. The group mean, by default, is the first regressor or covariate. In the primary study, BDI scores, age, sex are the next three regressors. Age and BDI scores were mean-centred (across all subjects) to enable the first regressor to be interpretable as the mean. In the follow up study, treatment (treatment received vs not treated) was included as the fifth regressor. To evaluate how regions in the network of interest interact, we used Bayesian model comparison to explore the space of possible hypotheses (or models). Candidate models were obtained by removing one or more connections to produce nested or reduced forms of the full model. As there is large number of possible nested models in the model space, the search algorithm used Bayesian model reduction (BMR) (41) that enables an efficient (greedy) search of the model space. BMR prunes connection parameters from the full model and scores each reduced model based on the log model-evidence or free energy. The process continues until there is no further improvement in model-evidence. The parameters of the selected models from this search procedure were then averaged, weighted by their model evidence (Bayesian Model Averaging) (116).

### Leave-one-out validation analysis

Finally, we tested whether the severity of depression could be predicted based on the modulation of effective connectivity. In other words, was the effect size large enough to have predictive validity. We chose connections that survived a threshold of 95 % posterior probability (very strong evidence) in the previous analysis (primary study). We used a leave-one-out scheme as described in (41). A parametric empirical Bayesian model was estimated while leaving out a subject, and was used to predict the BDI score of the left out subject, based on the specific connections chosen. The Pearson’s correlation between the predicted score and known score was calculated.

## Data and code availability

Our analysis code is available on GitHub (https://github.com/dipanjan-neuroscience/depression2021). Imaging data are available on Open-Neuro (https://openneuro.org/datasets/ds002748/versions/1.0.3 & https://openneuro.org/datasets/ds003007/versions/1.0.0).

## ACKNOWLEDGMENTS

We thank Prof. Mark Shtark who supervised the data collection and Dr. Andrey Savelov who conducted MRI and fMRI acquisition. This research was supported by the Basque Government through the BERC 2018-2021 program, by the Spanish Ministry of Science, Innovation, and Universities (BCBL Severo Ochoa excellence accreditation SEV-2015-0490 and BCAM Severo Ochoa accreditation SEV-2017-0718) and the project MTM2017-82379-R (AEI/FEDER,UE) (principal investigator: Dr. Maria Xose Rodriguez, BCAM). Data collection was funded by the Russian Science Foundation grant #16-15-00183. K.J.F. was funded by a Wellcome Trust Principal Research Fellowship 088130/Z/09/Z).

