## Supplementary figures for "Altered effective connectivity in sensorimotor cortices: a novel signature of severity and clinical course in depression"

### Group Mean Effective Connectivity

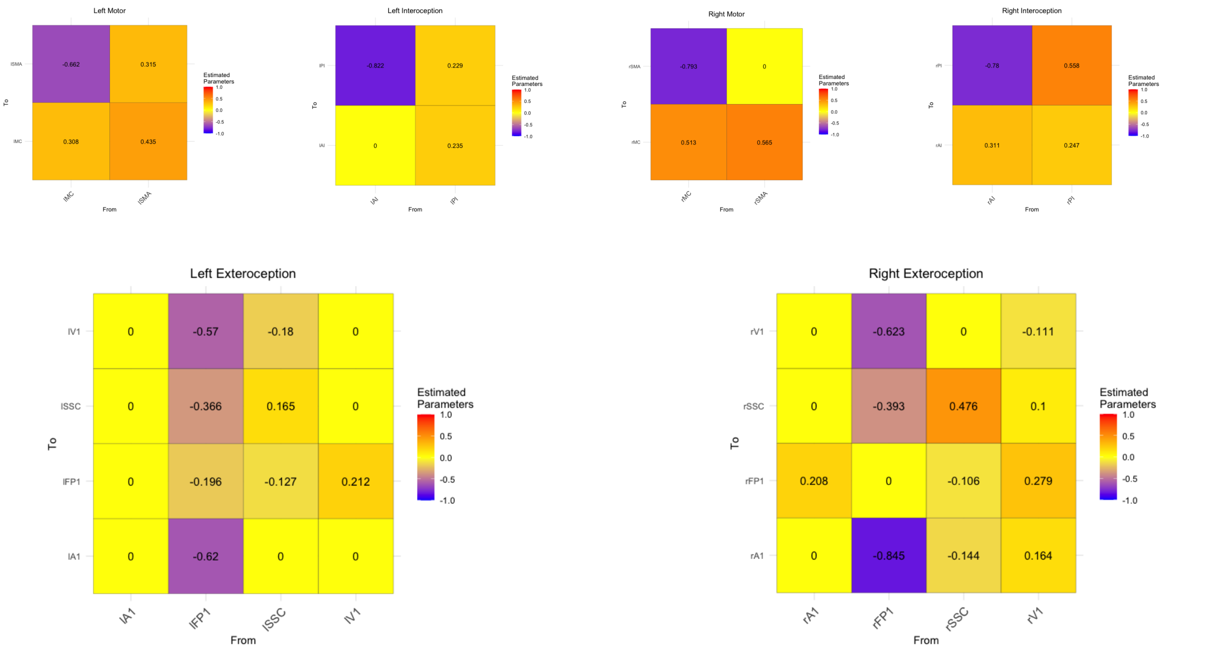


### Changes in effective connectivity with BDI scores

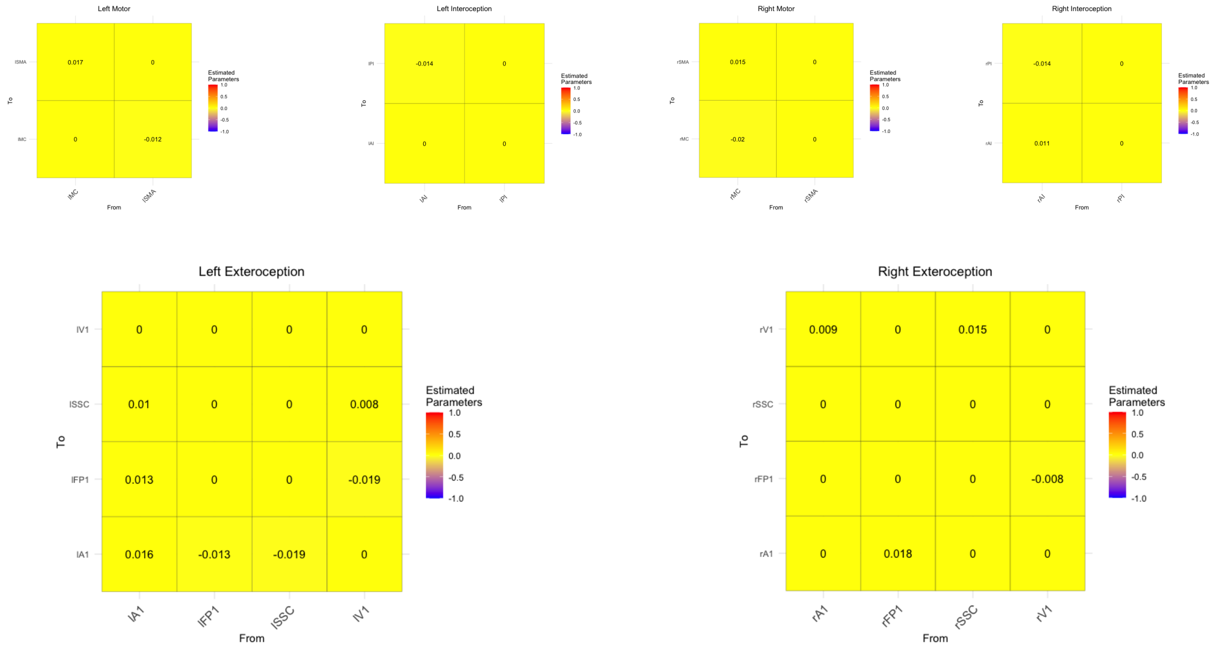


Supplementary Figure 1: Estimated parameters in the primary session

### Group Mean Effective Connectivity

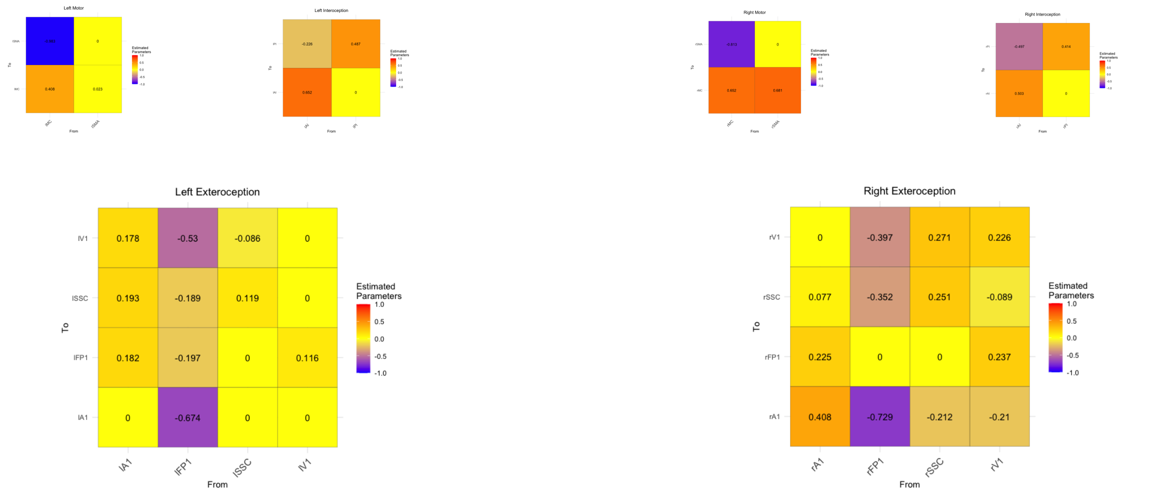


### Changes in effective connectivity with BDI scores

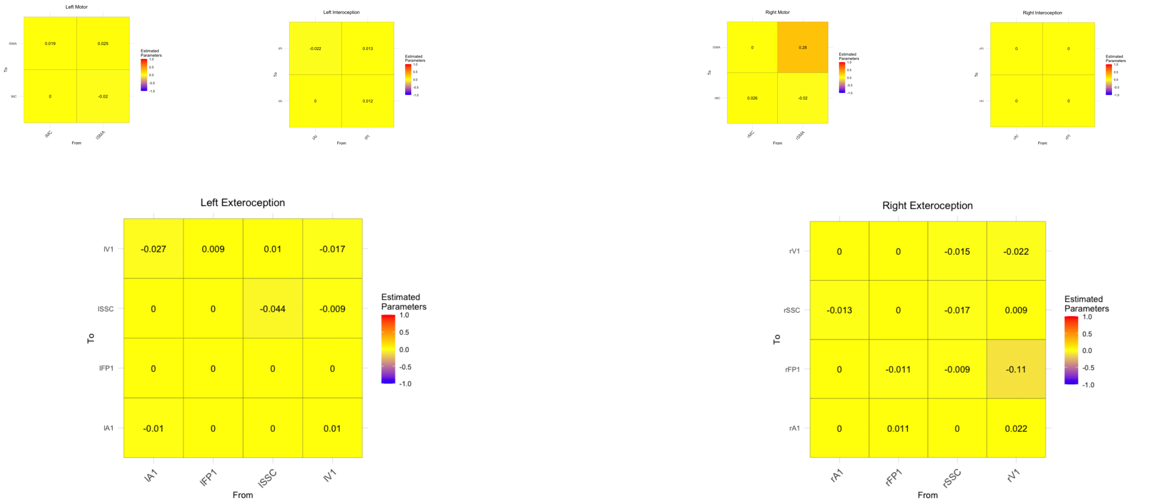


### Changes in effective connectivity with treatment

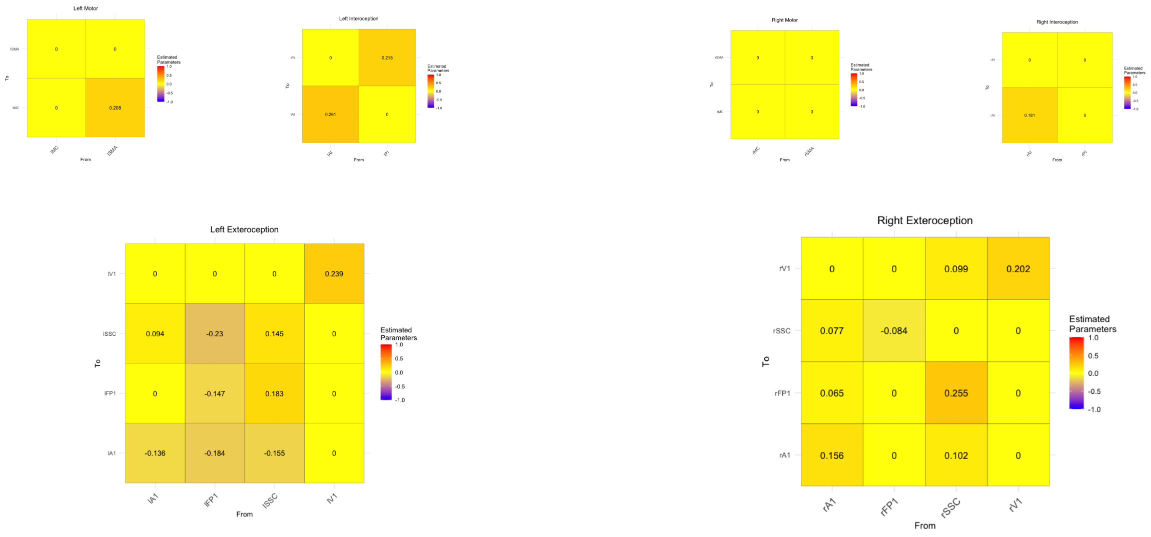


Supplementary Figure 2: Estimated parameters in the follow-up session
